# Genetic sequences are two-dimensional

**DOI:** 10.1101/299867

**Authors:** Albert J Erives

**Author notes:** **Abbreviations** 1-D: one-dimensional 2-D: 2-dimensional CDS: protein-coding sequence GA: gapped alignment indels: insertion and deletions MGMA: minimally-gapped MHA aligner MHA: maximal homology alignment MSA: multiple sequence alignment MSR: micro-satellite repeats NEE: neurogenic ectoderm enhancer TFBS: transcription factor binding site TR: tandem repeats *vnd*: *ventral nervous system defective* WCR: width cinch ratio.

## Abstract

In attempting to align divergent homologs of a conserved developmental enhancer, a flaw in the homology concept embedded in gapped alignment (GA) was discovered. To correct this flaw, we developed a methodological approach called maximal homology alignment (MHA). The goal of MHA is to rescue internal microparalogy of biological sequences rather than to insert a pattern of gaps (null characters), which transform homologous sequences into strings of uniform size (1-dimensional lengths). The core operation in MHA is the “cinch”, whereby inferred tandem microparalogy is represented in multiple rows across the same span of alignment columns. Thus, MHAs have a second (vertical) paralogy dimension, which re-categorizes most indel mutations as replication slippage and attenuates the indel problem. Furthermore, internally-cinched, inferred microparalogy in a self-MHA can later be relaxed to restore uniformity to 2-dimensional widths in a multiple sequence alignment. This de-cinching operation is used as a first resort before artificial null characters are used. We implement MHA in a program called *maximal*, which is composed of a series of modules for cinching and cyclelizing divergent tandem repeats. In conclusion, we find that the MHA approach is of higher utility than GA in non-protein-coding regulatory sequences, which are unconstrained by codon-based reading frames and are enriched in dense microparalogical content.

## Introduction

We describe a new approach to biological sequence alignment following attempts to perform gapped alignment (GA) starting at the poorly-conserved edge of a conserved developmental transcriptional enhancer. This new approach is based on a re-evaluation of evolutionary homology at small length-scales (“site positional homology”) and how it is or is not encapsulated in the GA approach. We find that the modeling of homology in the GA approach is an approximation that works best in protein domain-encoding sequences, which are doubly constrained by a codon-triplet reading frame and their encoding of secondary structural elements. In contrast, the modeling of homology in the GA approach is a poor approximation in *cis*-regulatory sequences, which are not so constrained (*e.g*., see Brittain *et al*., 2014).

In evolutionary genetics, sequence homology at a single site refers to identity of both position and symbol (letter). Two unlinked nucleotide sites located in two related sequences are considered to be homologous to each other based on **(*i*)** a globally-influenced inference of homology of position, and then **(*ii*)** a confirmatory inference of symbol identity at that homologous position. Thus, non-identical letters in a single alignment column of a gapped alignment still retain homology of position.

GA is based on a simplification of the homology concept as strict 1-to-1 orthology or lack thereof. If a sequence in an alignment is found to lack a symbol at an inferred alignment position, GA algorithms insert a null symbol represented by the dash character (“-”). The dash symbol does not carry information about its temporal polarization, *i.e*., whether it represents an insertion or a deletion in one sequence relative to another. The null symbol thus serves to adjust the non-uniform lengths of sequences or windows of sequence. With maximal homology alignment (MHA), a method introduced here, we use a robust and theoretically-correct alternative to the non-uniform length problem solved by GA’s null character insertions.

In the formalism of evolutionary genetics, homology is composed of two complementary categories known as orthology and paralogy. The orthology concept is meant to describe relationships connected through a single common ancestor, while the paralogy concept is meant to compartmentalize relationships complicated by duplications in one or more lineages of interest. In the current comparative genomics era, where the ancestry of gene families can be more comprehensively evaluated, the definitions of orthology and paralogy have been adjusted to maintain clade-level utility. For example, two related organisms can be defined to have orthologous genes if these genes are descended from a single gene present in their common ancestor, regardless of lineage-specific duplications since this common ancestor (Remm et al., 2001; Hubbard *et al*., 2007). This particular definition is computationally-compatible with the need to distinguish ancient duplications preceding the common ancestor (the “out-paralogs” that define large gene super families) from recent lineage-specific duplications (“in-paralogs”), which are often found in one lineage or another (Remm et al., 2001). We now illustrate a fundamental flaw of GA by analogy to the phylogenetic analysis of gene families with their many gene duplications.

If gene *A* from organism “X” is being aligned to a duplicated pair of genes *A1* and *A2* from organism “Y”, this alignment will by composed of three rows of gene sequences (*A*, *A1*, and *A2*) and *n* alignment columns, as determined by the lengths of the sequences and the number of dashes invoked by the chosen GA method. For multiple sequence alignment (MSA) of the gene family repertoires from different genomes, incorporation of paralogous genes from the same genome is readily permissible, and is informative for distinguishing the ancient paralogy groups (*i.e*., ancient duplications) of a large superfamily from the lineage-specific duplications within one paralogy group. In fact, MSA can proceed as if all the ancestral and lineage-specific gene duplications present in any one genome came from different genomes. Thus, most phylogenetic analyses treat the set of genes *A*_X_, *A1*_Y_, and *A2*_Y_ from two different organisms in the same way as a set of three orthologous genes *A*_X_, *A*_Y_, *A*_Z_ from three different organisms.

In GA, the categorical partition of homology into orthology and paralogy is not incorporated in the sense that it does not include “microparalogy” of position, by which we mean internal tandem duplications typical of replication slippage (<< 100 bp) (Strand et al., 1993). There are likely several reasons for why GA has not been built to handle microparalogy. First, while paralogous gene duplications are discrete and more easily recognized as separate entities, with small internal tandem repeats, and much more so with small unstable repeats of repeats, this useful discreteness of duplicated segments is more ambiguous. Second, microparalogical duplications are more numerous and present in a more uniform distribution. Third, it is not necessarily predetermined that the different lengths of homologous segments will often be due to replication slippage (Strand et al., 1993; Haber and Louis, 1998), and in fact other complex mechanisms have been proposed (*e.g*., Amos, 2010). Fourth, GA methodology has matured over the decades on an example corpus enriched in the amino acid sequences of protein-coding domains with well-defined secondary and tertiary structures, as well as the nucleotide sequences encoding them.

With GA, if replication slippage produces a repetition of the heptamer 5´-CAGGTAG into 5´-CAGGTAG<u>CAGGTAG</u>, one of the two halves of the 14 bp sequence will be aligned to dashes inserted in other sequences. Nonetheless, positions 1 and 8 in the duplicated sequence are homologous to each other via local microparalogy, and furthermore both positions 1 and 8 are equally homologous to position 1 in the unduplicated homologs. Thus, any *single* multiple sequence alignment (MSA) produced by GA disperses a great deal of local microparalogy into separate alignment columns, causing the need to insert null characters in other sequences. This is an issue exacerbated by the immense force of replication slippage in sequence evolution particularly in *cis*-regulatory sequences (Crocker et al., 2010; Gemayel *et al*., 2010; Kelkar *et al*., 2011; Ananda, *et al*., 2013; Duitama *et al*., 2014; Brittain, et al., 2014).

Under GA, the magnitude of the so-called “indel problem” has propelled an almost 50 year-long search for a “master equation” solution, which would provide an efficient and satisfactory mitigation of indels by dictating an optimal pattern of gap insertions (for example treatments spanning this time frame, see Sankoff, 1972, and Holmes, 2017). Even probabilistic approaches, which are amenable to relaxing the assumption of a single optimal or perfect gapped alignment and dealing with probability distributions of alignments, are nonetheless built on a GA framework (*e.g*., Satija et al., 2008). All of GA features a deeply-embedded world model involving computational decisions to match, mismatch, insert, and/or delete in order to restore a non-biological 1-to-1 versus 1-to-none concept of homology that is really positional orthology *sensu stricto* (see Table 1).

**Table 1.**
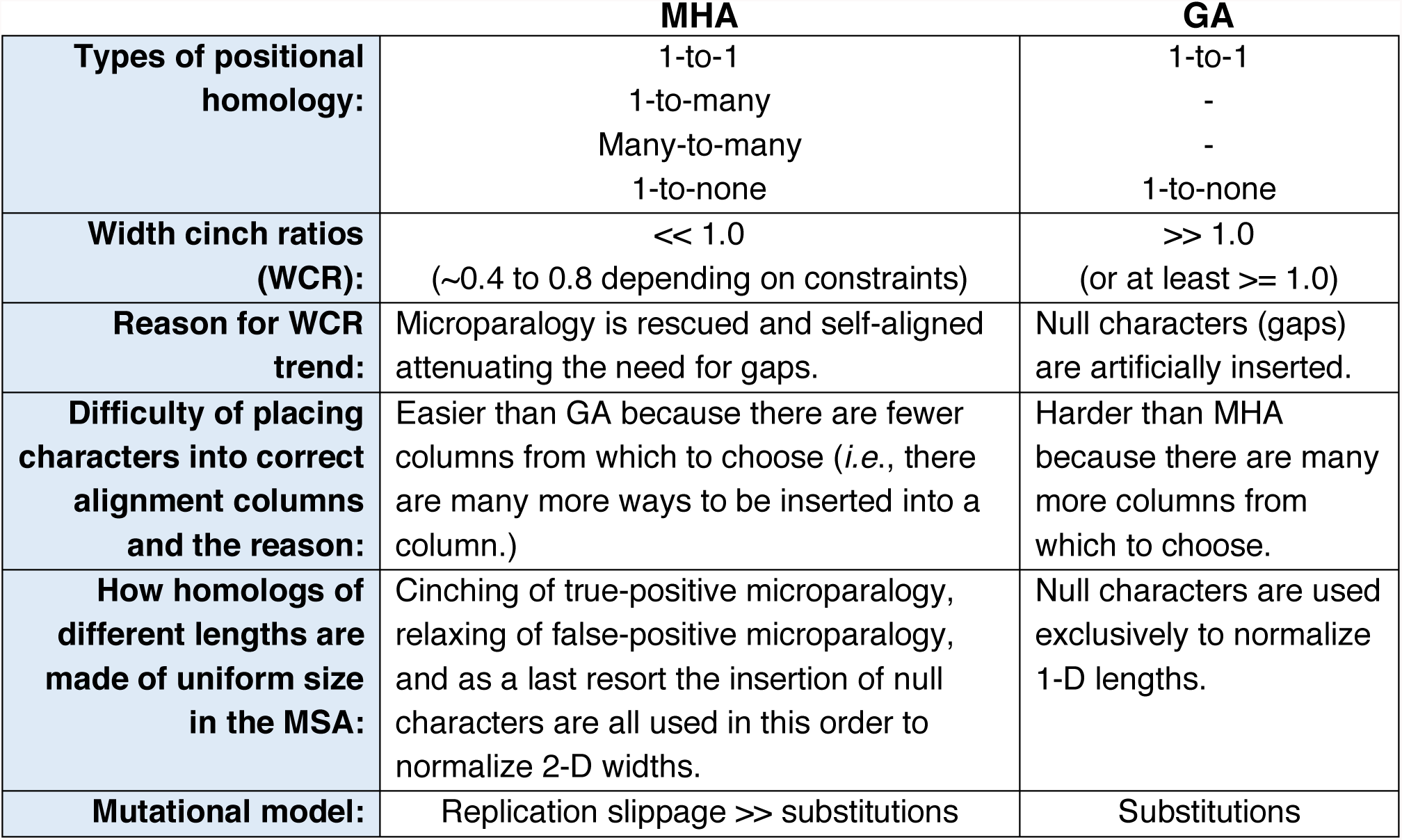
Summary comparison of MHA versus GA for MSA

Here, we define **maximal homology alignment** (**MHA**) as any method that models DNA sequence as a two-dimensional string of symbols according to a homology formalism that includes microparalogy (1-to-many and many-to-many correspondences, see Table 1). We first describe how the MHA method facilitates alignment of a highly-conserved transcriptional enhancer and compare these results with alignment of a protein coding sequence. We find that any GA approach is intractable in ways that are not an issue to MHA. We then implement a computational solution to MHA and demonstrate its utility. We distinguish between MHA as a general goal from our specific implementation in a program called *maximal*. The *maximal* implementation may represent one of several ways to achieving MHA and so we refrain from making statements about its relative computational efficiency in the space of MHA solutions. Nonetheless, in our sparse matrix implementation, the initial MHA preparation of a single sequence of length *n* has a time complexity that is linear in *n*, *i.e.*, with a run-time of *O*(*n*), and produces strings with a radically diminished need for computationally-intense gap solutions during MSA. In addition to the attenuation of length non-uniformity of homologs, a multiple MHA aligner can draw upon internally cinched characters as a first resort and null (gap) characters as a last resort. Thus, MHA has a more powerful and biologically-relevant basis for the restoration of length uniformity to a set of homologs in a multiple sequence alignment.

## Results

### Alignment at the edge of a transcriptional enhancer

We demonstrate the need for the MHA approach with an example sequence beginning at the poorly-conserved edge of a highly-conserved intronic transcriptional enhancer (Erives and Levine, 2004; Crocker and Erives, 2013). This enhancer occupies the second intron of the *Drosophila* gene *ventral nervous system defective* (*vnd*), which is homologous to the vertebrate *NKX2-2* and *NKX2-8* genes. The *vnd* intron harboring its neurogenic ectoderm enhancer (NEE) does not contain any other enhancer activity or detectable sequence conservation besides the regulatory belt of peaks constituting the binding sites within the enhancer (Fig. 1a). In previous studies, we have documented the evolutionary role replication slippage plays in shaping these enhancers (Crocker et al., 2010; Brittain, et al., 2014), similar to processes shaping the human genome (Gemayel *et al*., 2010; Kelkar *et al*., 2011; Ananda, *et al*., 2013; Duitama *et al*., 2014).

**Fig. 1.**
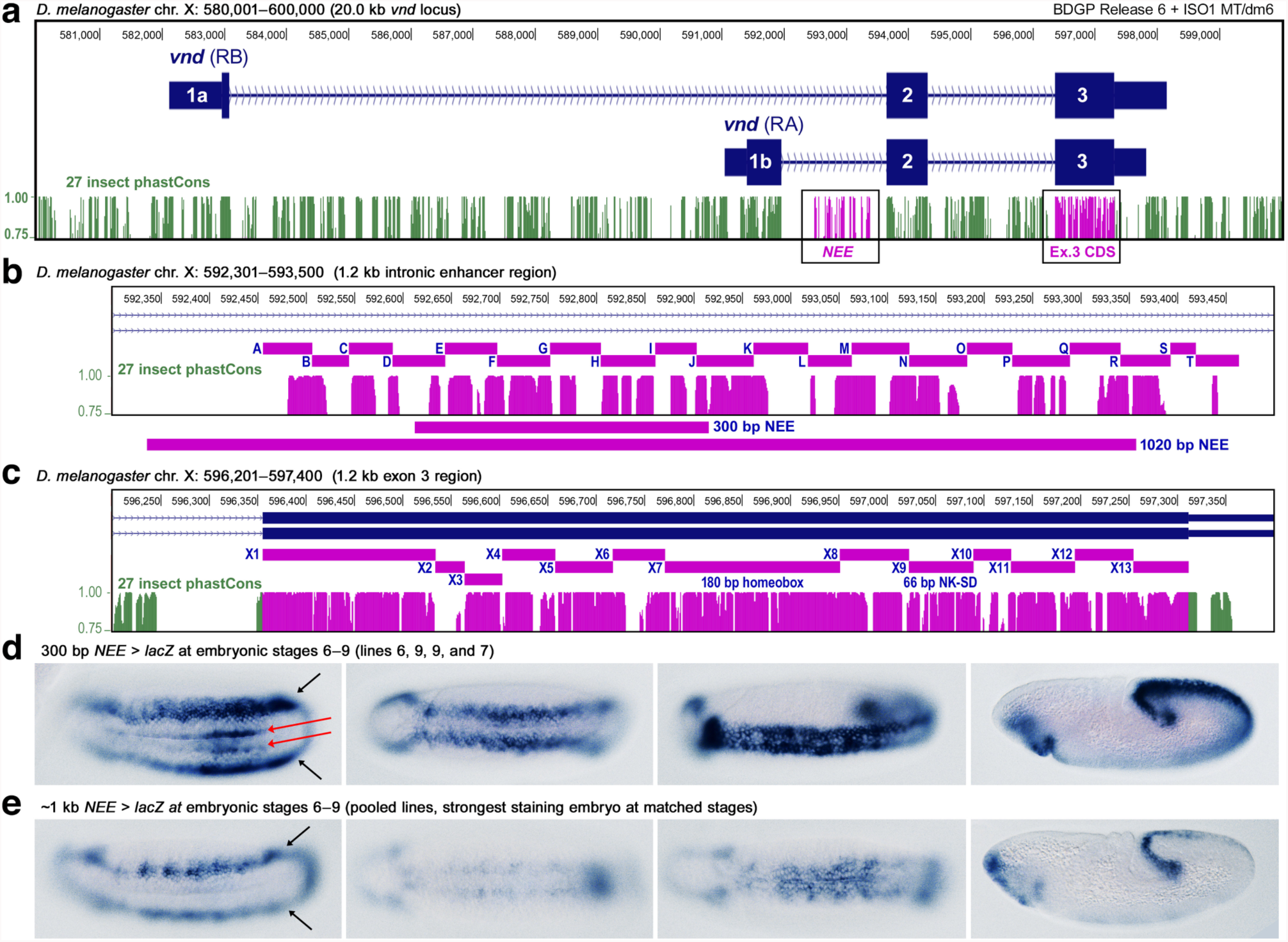
The full-length *vnd* neurogenic ectoderm enhancer (NEE) of *Drosophila* works as a unit for correct spatial and temporal patterning. This study introduces maximal homology alignment (MHA) to study and compare the evolution of an enhancer DNA sequence and an adjacent protein-coding sequence from the *Drosophila* locus *ventral nervous system defective* (*vnd*), a highly conserved homolog of vertebrate *NKX2.2*/*2.8*. **(a)** Depicted is the 20 kb *vnd* locus from *Drosophila melanogaster* showing the 27 insect phastCons conservation score (bottom track in green and magenta). Two 1.2 kb regions containing the <u>n</u>eurogenic <u>e</u>ctoderm <u>e</u>nhancer (NEE) and the exon 3 protein-<u>c</u>oding <u>D</u>NA <u>s</u>equence (CDS) are both sites of microfoam evolution, which occurs in the interstices between adjacent conserved elements. **(b)** The 1.2 kb NEE was divided into 20 blocks that are alignable in closely related species (blocks lettered A–T). This region encompasses the 300 bp minimalized NEE, which recapitulates the neuroectodermal expression at embryonic Stage 5 of both the endogenous locus and the longer NEE reporter. **(c)** The 1.2 kb region of the terminal exon, which contains both the 180 bp homeobox (X7 segment) and the 66 bp encoding the NK-specific domain (X9 segment), was divided into 13 alignable segments numbered X1–X13 as shown. **(d)** Shown are transgenic embryos stained with an anti-sense RNA *lacZ* probe to detect robust lateral neuroectodermal (black arrows) and ectopic mesodermal (red arrows) beginning at embryonic stage 6 and continuing until the long germ-band extended stage 9 embryo (rightmost panel). **(e)** Shown are embryos matched by stage and orientation to those immediately above in panel D. Unlike transgenic embryos carrying the minimalized 300 bp NEE, the ~1 kb NEE reporter drives limited expression (fewer embryos and less robust expression) in embryonic stages after stage 5, likely due to conserved arrays of homeodomain binding sites (see text and SI).

To perform a fine-grained sequence alignment of the *vnd* NEE, we segmented this enhancer into 20 lettered sections, beginning with the 51 bp segment of Block A (see Fig. 1b, and Fig. 2a). Figure 2b shows a state-of-the-art Multiz Alignment based on the sequences from 27 insects (Blanchette *et al*., 2004). Although this example highlights the specific output from Multiz, the points that we will make are general to gapped alignment. In this alignment we highlight several problems relative the maximal homology alignment. For now, we merely point out that not all of these 9 shown species begin with sequence as three species begin with dashes (“-“) and another begins with unalignable sequence (“=”). As we shall see, both of these issues can be corrected with MHA.

**Fig. 2.**
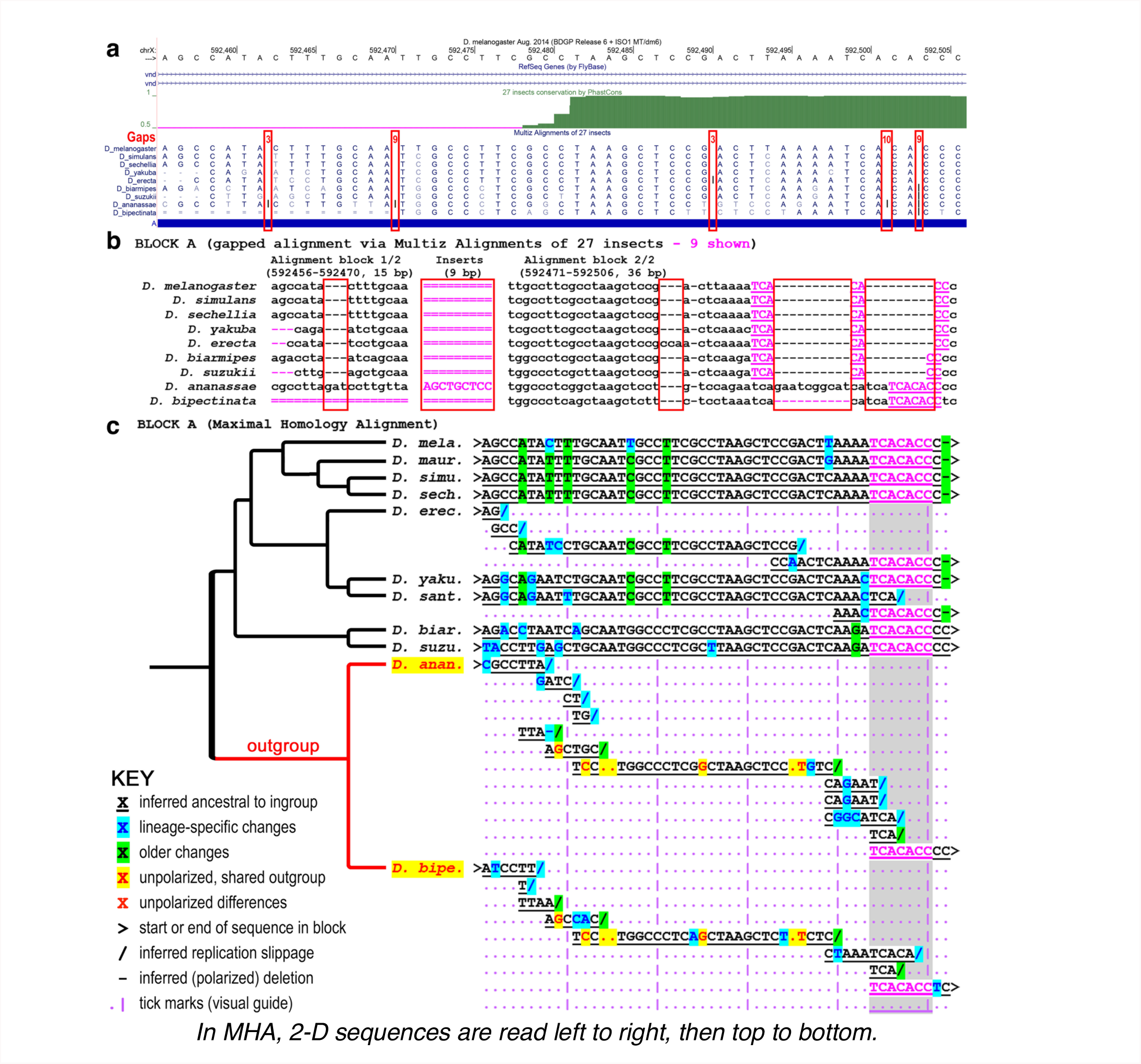
Maximal homology alignment (MHA) rescues orphaned microparalogy. Depicted is the alignment of block A of the *vnd* NEE region in a standard genome browser view **(a)**, and expanded multiple sequence alignment (MSA) based on the 27 insect Multiz pipeline **(b)**, and MHA **(C)**. Block A is the beginning of detectable homology at the edge of a highly conserved developmental enhancer. **(a)** Genome browser views are necessarily relative to the assembly of a reference genome, which is likely to have sequences missing relative to related genomes. These are known as gaps in the alignment and can be graphically annotated as shown without disrupting the intactness of the reference sequence. **(b)** The expanded 27 insect Multiz alignment is typical grossly distorted by the inclusion of many gaps. **(c)** MHA rescues the vast majority of insertions in specific lineages as microparalogy. This type of MSA makes it easier to read and detect intact TFBS-like sequences that are conserved in each sequence. In gapped alignment these conserved TFBS-like sequences are mistakenly aligned to specific duplicated fragments in an attempt to minimize the insertions of gaps. Gapped MSA is thus fatally flawed by not incorporating microparalogy. Cladogram is intended only to show species relationships.

In general, gapped alignment includes many dashes (Fig. 2a). These null characters represent that there are either insertions or deletions relative to one another. However, as to whether insertions or deletions predominate is not immediately clear. Additional sequences are present in one sequence that are not alignable to others.

The heptapeptide sequence 5′-TCACACC is present in each of the *Drosophila* species (highlighted in magenta in Fig. 2), but the gapped alignment splits this sequence to match various sub-strings present within the a large gap (compare Fig. 2b to Fig. 2c). Some of these fragmentary alignments, such as the central 5′-CA (nucleotides 4 and 5) occurs due to a gap parsimony-constraint in the MSA (highlighted gap).

Fig. 2c shows the hand-aligned MHA version of the multiple sequence alignment of the first block, Block A, of the *vnd* NEE. In stark contrast to the many dashes in the gapped MSA, we find that there are only three nucleotides of unpolarized indels (highlighted dots) in the two outgroups sequences (red, yellow highlight). In addition, we infer that there has occurred at most two high-confidence deletions: a single nucleotide contraction of the a run of 4 C’s in the ancestor of the melanogaster group, and potentially a second single letter contraction of a run of three A’s in *Drosophila ananassae*. Last, the vast majority of indels marked in the gapped 27-Multiz alignment is explained by replication slippage. Furthermore, we identify that the central problem of all regulatory alignment is the evolution of private, which is to say lineage-specific, microfoam sequence. This microfoam sequence can be explained as divergent replication-slippage involving repeats of repeats.

To demonstrate the appeal of a possible MHA strategy, we look at the first ~50 bp of block-A for just a pair of flies. An example of pair-wise GA (Needleman-Wunsch) for this example non-coding sequence is shown in Fig. 3a. These poorly conserved sequences are 46 bp long in *Drosophila melanogaster* and 51 bp long in *Drosophila erecta* and have been diverging for at least 10 million years. This sequence is the first detectable window of alignable sequence on the edge of this enhancer and can be truncated without loss of correct spatial patterning of a reporter gene. Because the sequences are of different lengths, gaps are inferred in multiple places (Fig. 3a).

**Fig. 3.**
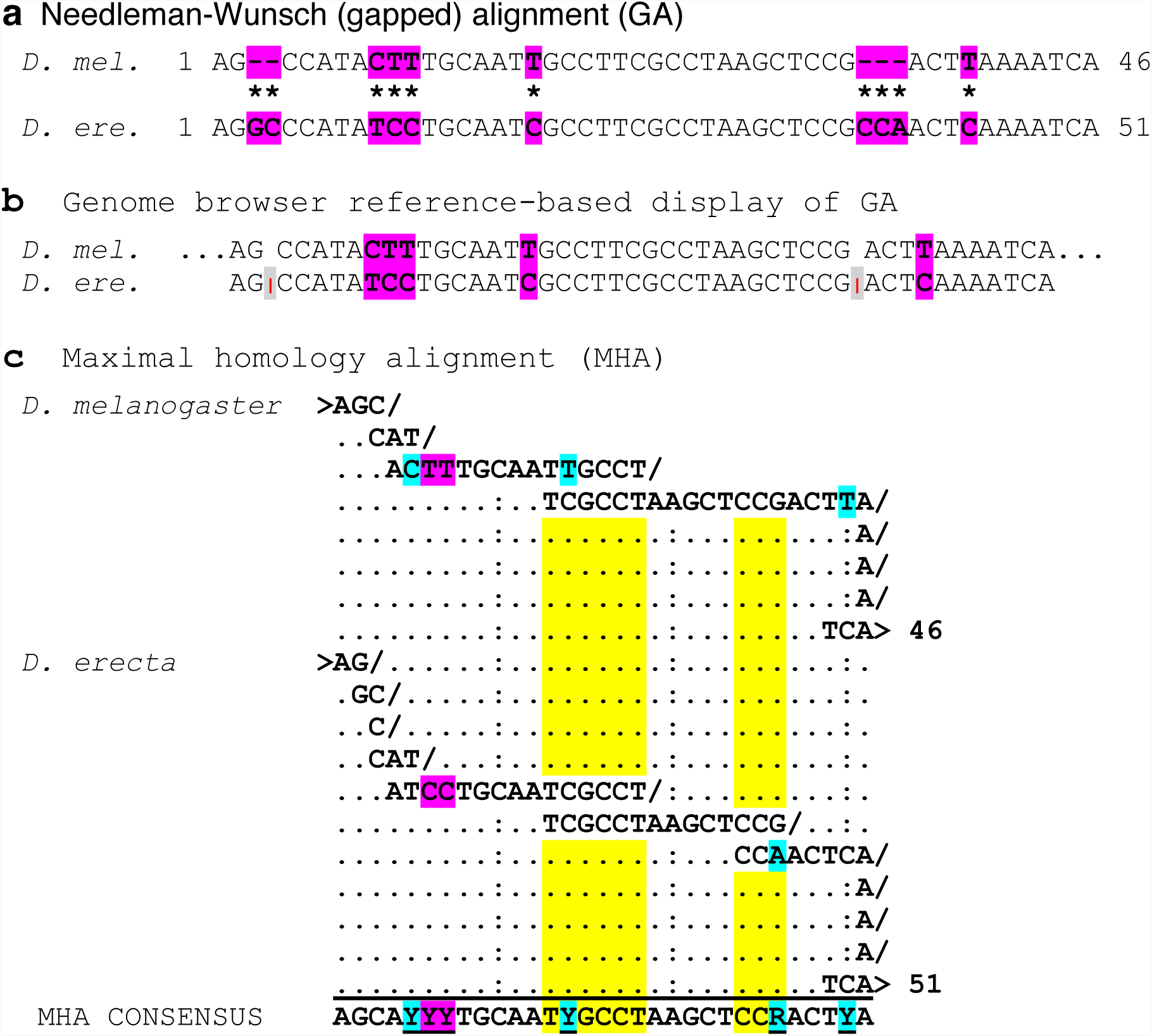
Self-MHA at the edge of a transcriptional enhancer can produce automatic alignment. MHA dispenses with the need to solve an artifactual indel problem that is magnified in GA. **(a)** Example of GA using the global alignment method of Needleman-Wunsch. Shown are just the first ~50 bp of the window of sequence shown in Fig. 2 for only a pair of *Drosophila* species. The differences are all highlighted in magenta to represent that these are unpolarized in the absence of outgroup sequences, which would help determine the ancestral characters. **(b)** Example of the same window of sequence in a genome browser view from the perspective of the reference species *Drosophila melanogaster*. Sequences present only in *D. erecta* are hidden. **(c)** Self-MHA preparation of the pair of enhancer edges is sufficient to bring the divergent sequences into alignment. The cyan highlighted sites are polarized substitutions based on the pair-wise alignment and later confirmed by consulting outgroups. The sequences proceed left to right, top to bottom and forward slashes (“/”) are used to indicate inferred replication slips. The “>” symbol is used to mark the start and end of each species’ sequence. Two regions representing an ancestral *k* = 6 TR and a more recent lineage-specific *k* = 3 TR are highlighted (yellow). These replication slips are not immediately obvious in GA representations. Remarkably, each highlighted region allows polarization of one substitution each (highlighted in cyan) without the use of a third outgroup sequence. Tighter cinching of the pyrimidines at positions 5–7 could have also have been conducted but is not shown here for clarity.

In a genome-browser graphical representation of the same pair of aligned sequences, an additional constraint on displaying sequence homology can be demonstrated (Fig. 3b). In a genome browser view of the same pair of sequences, one displays homologous sequences only if they have been inferred to have a 1-to-1 GA relationship to the reference genome positions. Thus, extra sequences in other related genomes are typically hidden, at least at the level of a zoomed-in genome browser-like display (tick marks in Fig. 3b). This design choice is a necessary constraint imposed by GA in general because different related genomic sequences will have private insertions in different places, thus making visual display to the reference unwieldy.

In contrast to GA, MHA allows one to include microparalogy in the alignment (Fig. 3c) that is otherwise hidden either by obscurity (Fig. 3a) or by constraint of design (Fig. 3b). Remarkably, this pair-wise MHA reveals that self-MHA preparation has done most of the alignment work without any direct pair-wise comparisons. Furthermore, MHA also allows the inference of polarity in this pair-wise alignment without the use of a third sequence in a known outgroup relationship. Indeed, with multiple repeats, polarity can be inferred with only a single sequence.

This work was heuristically guided by the complete hand-MHA version of the *vnd* NEE and *vnd* exon 3 for comparison (Supporting File S1). Example blocks from this MHA are shown in Figures 4 and 5, which demonstrate the utility of MHA for both *cis*-regulatory and protein-coding sequences. We thus decided to explore this method further by developing software tools to conduct MHA. Before describing the specific software programs we have produced, we first address some questions about how MHA would work in comparison to GA.

**Fig. 4.**
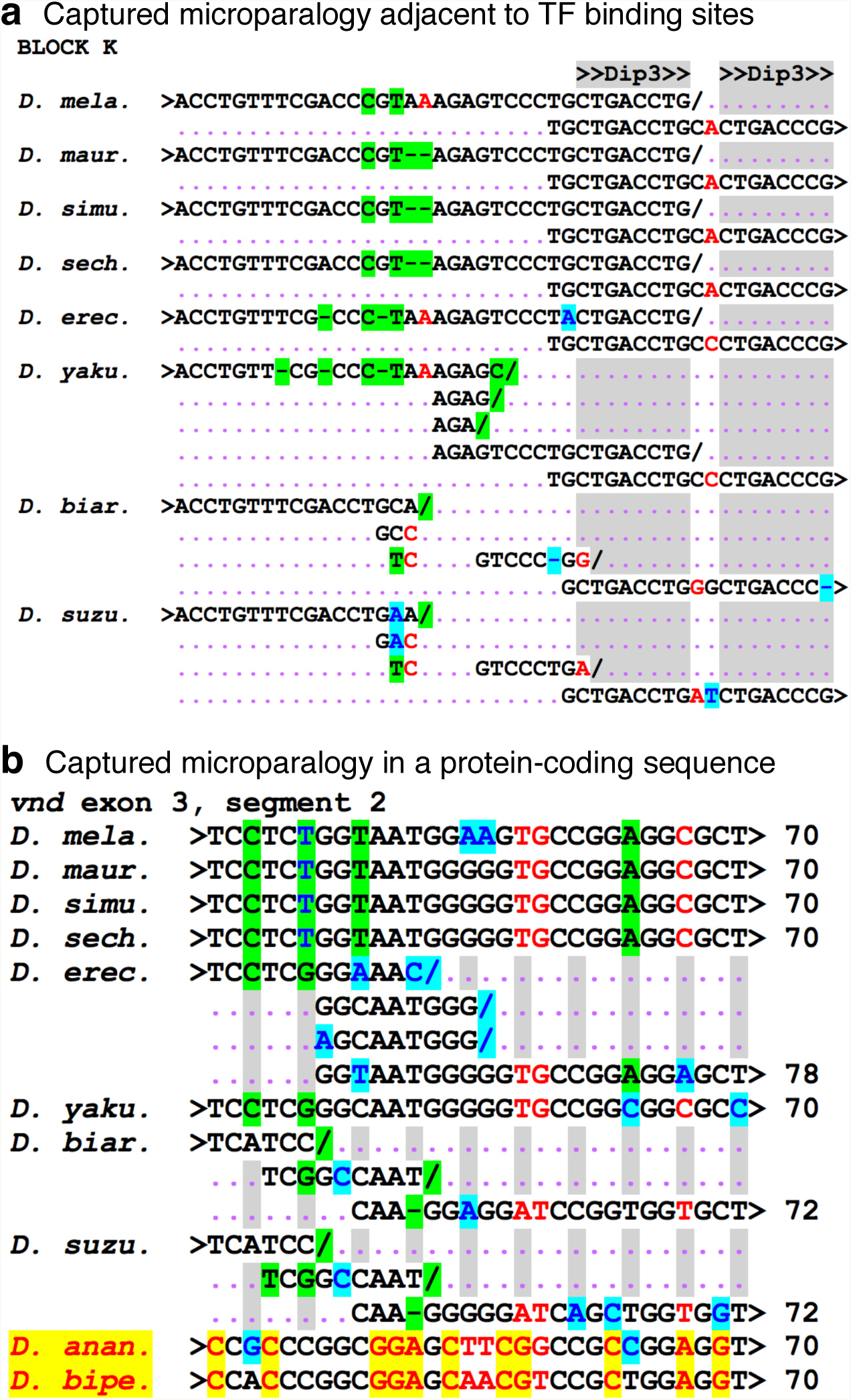
Microparalogy accumulates in *cis*-regulatory and protein-coding sequences. Gapped alignment (GA) **(a)** Example of imperfect microparology adjacent to and within a cluster of Dip-3 sites in the vnd NEE. Also note that the array of Dip3 sites also constitute a type of imperfect tandem repeat that is evolving. **(b)** For comparison is shown some protein-coding sequence from the same locus. This region occurs upstream of the adjacent homeodomain (HD) and NK-2 specific domain, which do not allow local microparalogy to accumulate. The first gray-shaded column on the left marks the third position of the first codon but due to the MHA, it does not always coincide with the third position after certain slips. This demonstrates that indels in protein-coding sequence are sometimes compensated by in frame indels elsewhere.

**Fig. 5.**
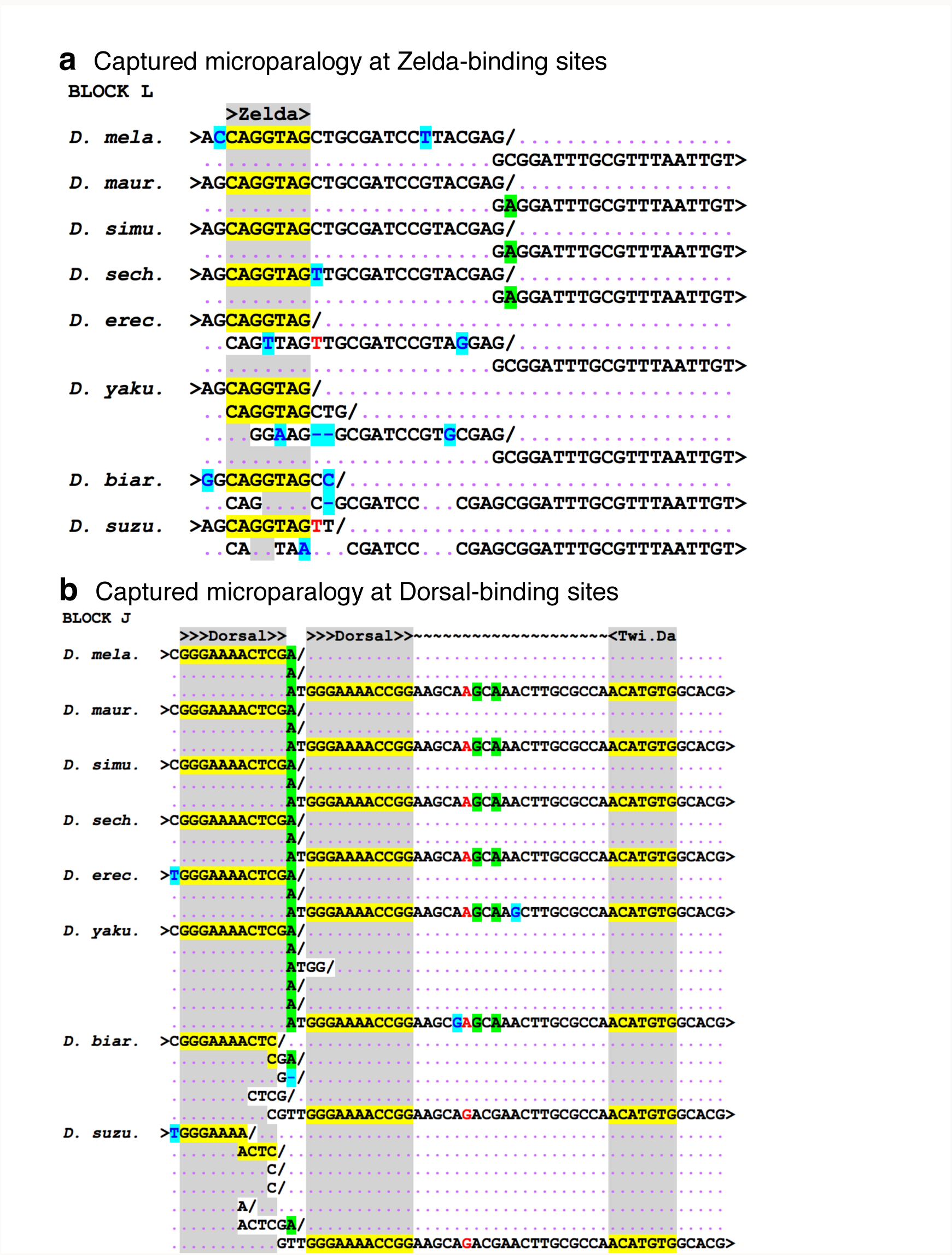
TF binding site turnover via microparalogy. **(a)** Example of the hand-aligned MHA at a Zelda site. This region is chopy but most of the alignment can be accomplished with just a few deletion characters (en dashes). This MHA can thus be accomplished without null character being inserted. **(b)** Microparalogy of a Dorsal binding site in the MHA shown is consistent with our previous conclusion that many of the Dorsal binding sites are relics produced by selection for alternative spacer lengths relative to the coordinating Twi:Da E-box binding site (shaded site on the right). In the *D. suzukii* sequence (bottom-most sequence) the pentameric sequence 5´-(AACTC) repeats twice with the initial A’s and final C of the first unit continuing longer homolomeric runs.

### How MHA works in comparison to GA

One important question concerning MHA is how one should evaluate whether a genetic sequence is cinched-well or even how much one should cinch internal homology. In principle, some cinched sequence will represent true-positive microparalogy that has been restored to co-occupy a set of alignment columns. However, some cinched sequence may represent false-positive microparalogy of adjacent sequences that have become similar via mutational drift. Thus, the more aggressive one cinches microparalogy, for example by including the capacity for modeling repeat divergence, the likelier it is that false-positive microparalogy will be cinched.

False-positive microparalogy is not likely to be an issue to MHA from the following rationale confirmed by our initial exploration of the method. We can describe gapped alignment as the process by which artificial null characters are distributed into a set of homologous sequences in order to restore sequence alignment and to effectively make them of uniform lengths as part of a multiple sequence alignment (MSA) (Fig. 6a). In a similar way, we can envision MHA as producing an MSA that is composed of homologs of uniform 2-D widths (Fig. 6b), but with several key differences.

**Fig. 6.**
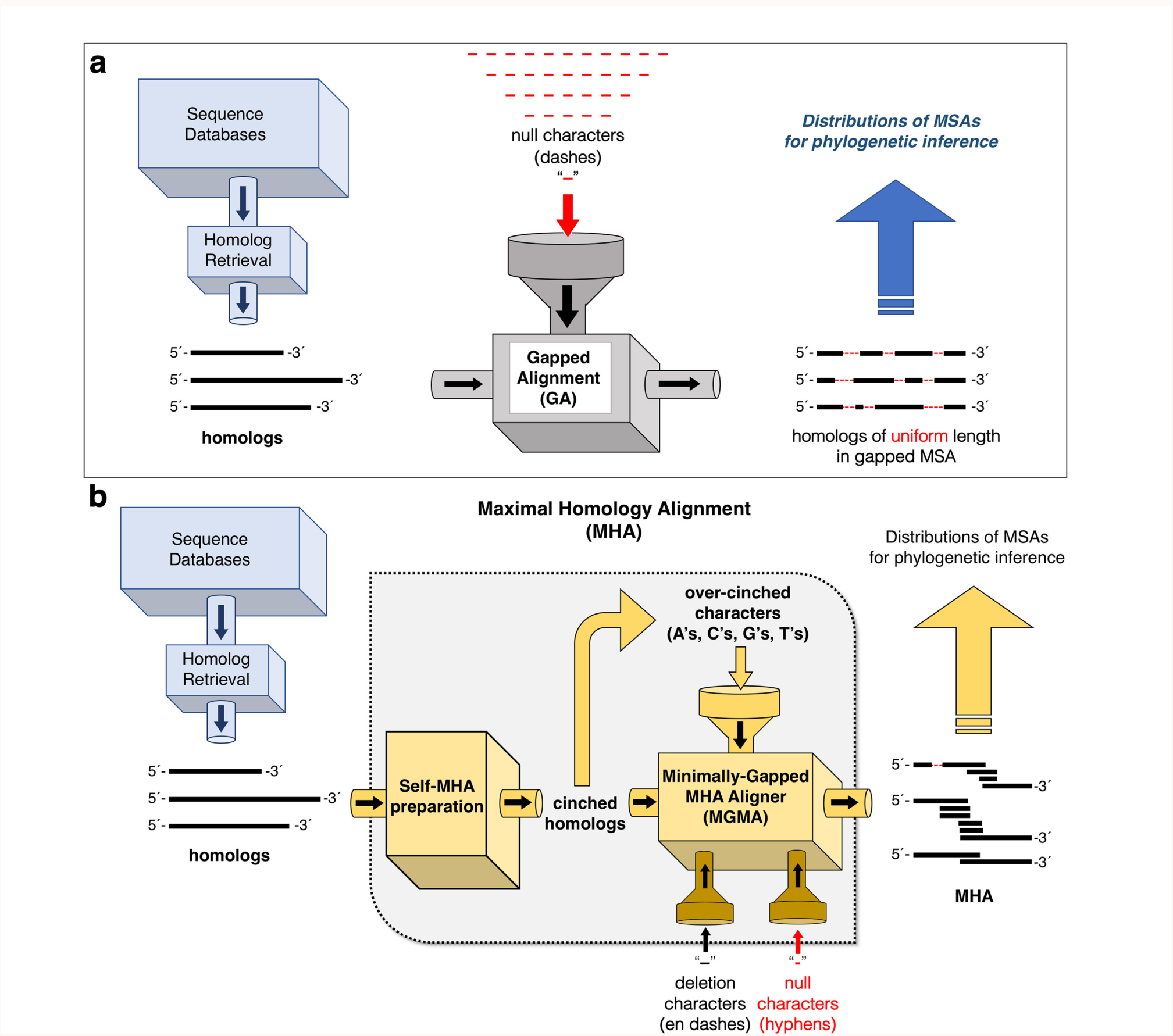
MHA has a more powerful capacity to adjust the non-uniform lengths of homologs than GA. Gapped alignment (GA) **(a)** and maximal homology alignment (MHA) **(b)** are contrasted by how they transform homologous sequences into aligned strings of uniform length. In both approaches, unspecified database retrieval methods identify homologs for phylogenetic analysis. **(a)** In GA, null characters represented by the dash character are distributed into the homologous sequences in order to restore columnar alignment in a multiple sequence alignment (MSA). **(b)** In contrast to GA, MHA first transforms homologs via self-MHA preparation (first gold box on the left). This process results in cinched homologous sequences whose internal microparalogy is self-aligned in a 2-D MHA and attenuates a major component of the indel problem. Cinched homologs are composed of correctly cinched true positive microparalogy and “over-cinched” non-paralogous sequence, which does not pose an issue to MHA. In the next step, cinched homologs are fed into an MHA aligner that works with 2-D self-MHAs to produce a minimally-gapped MSA of MHAs. If character insertions are required to restore sequence alignment, the MGMA module first attempts to use local cinched characters. If this is not possible, MGMA uses other characters. Unlike GA, the MHA approach can infer true deletions in regions of internal microparalogy. In this case, a deletion character is used in the specific sequence (the en-dash “–”). In other cases, and as a last resort when one of the homologs contains un-cinchable inserted sub-sequence, the null character, now represented by the hyphen (“-”) is used to restore alignment of all homologs lacking the inserted sub-sequence.

MHA can be conceptually broken into two separate procedures involving a self-MHA preparation module and a multiple-MHA aligner. Because the multiple-MHA aligner is designed to produce a minimally-gapped MSA we refer to it as minimally-gapped MHA aligner (MGMA). Thus, the first key difference between GA and MHA is that the former aligns 1-D sequences and the latter aligns 2-D sequences that were prepared by self-MHA. The former GA method produces MSAs of uniform 1-D lengths and the latter MHA method produces MSAs of uniform 2-D widths.

A second important difference is that GA works only to restore alignment via insertion of null characters (dashes). In contrast, MHA can draw upon the internal cinched characters present in the self-MHA to restore alignment. This powerful capacity to avoid using null (gap) characters is a fundamental departure from GA. Furthermore, the need to draw upon internal cinched sequences to restore homolog alignment is most likely to occur with false-positive microparalogy. To summarize, overcinched false-positive microparalogy is used as the first resort by the MGMA module rather than null characters from the void. This makes MHA a much more powerful approach for restoring uniform lengths to divergent homologs.

### Implementation of MHA in the *maximal* program of cinching modules

#### Path traversion

The *maximal* implementation of MHA begins similarly to the dynamic programming strategy used in global (*e.g*., Needleman and Wunsch, 1970) and local (*e.g*., Smith and Waterman, 1981) alignment with the construction of a path box, which when traversed according to the algorithm describes the cinching required to produce local self-alignment (Fig. 7a). The path box is filled with scoring values according to a substitution matrix. However, as substitution matrices are themselves based on MSA, the robustness of MHA over GA motivates a re-assessment of nucleotide and perhaps even amino acid substitution matrices. For example, in our initial explorations of MHA, we find there is a higher transition to transversion ratio for non-protein-coding regulatory sequences than has been previously reported, and this value is more in line with the higher ratios seen in protein-coding sequences (a more comprehensive assessment of this is being reserved for a later study).

**Fig. 7.**
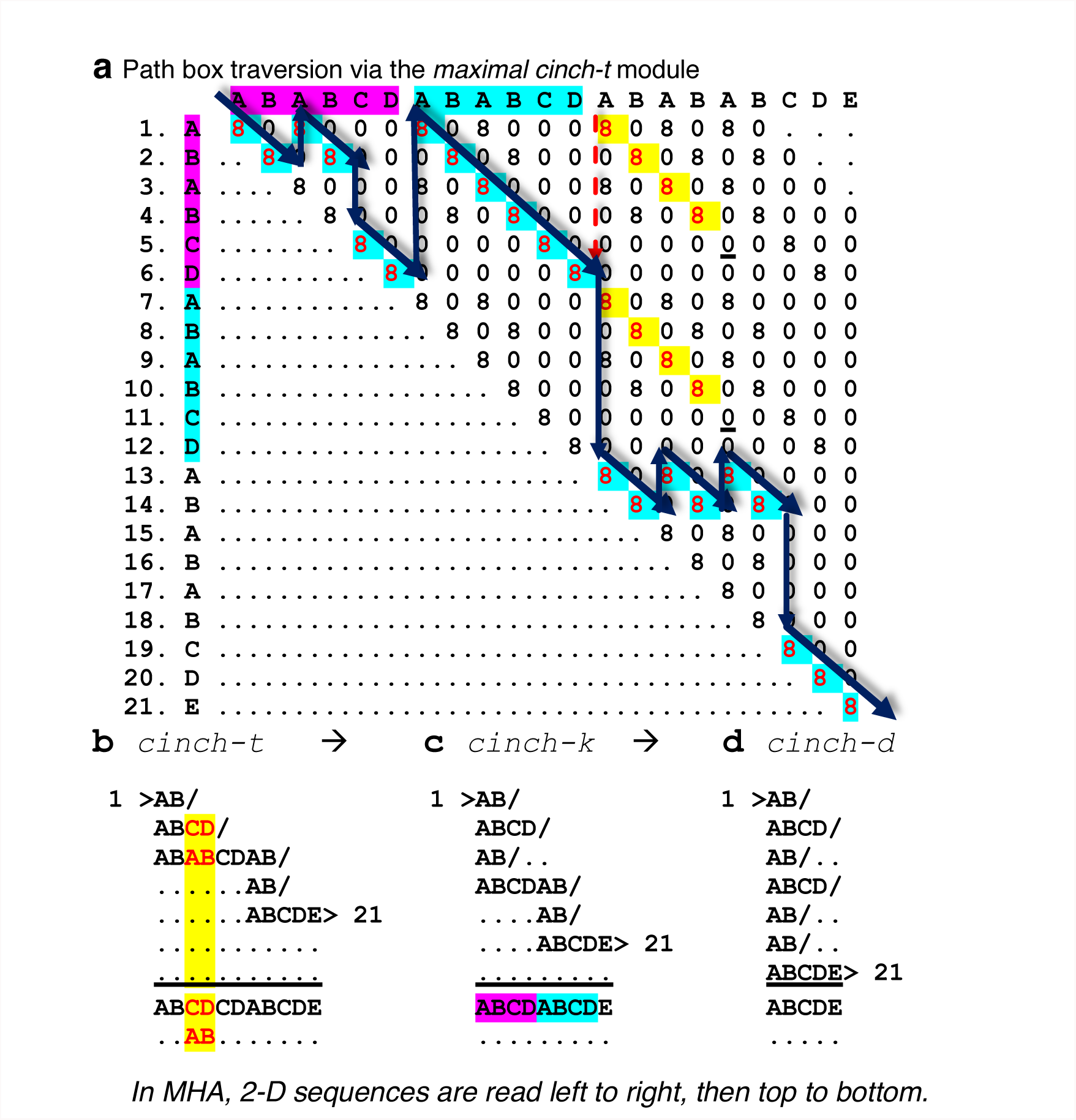
MHA is implemented in *maximal* via a series of cinching modules. **(a)** The *maximal* implementation of MHA begins with traversion of a hemi-diagonal sparse *m* × *n* matrix filled according to a substitution model in a self by self comparison. In this alpha-string example, the substitution model gives 8 points for a perfect match and 0 otherwise. Proceeding column by column left to right, positive scores in earlier rows are evaluated to see if they complete a diagonal of size *k* = *n – m* with a score above threshold. Identification of above-threshold diagonals constitute the initial path and are highlighted in cyan. Two diagonals highlighted in yellow are example diagonals that were disqualified for containing mismatches (underlined 0’s) before reaching the required *k*-mer length. **(b)** An initial *cinch-t* module produces a 2-D self-alignment based on the stipulated traversion of the pathbox that cinches tandem repeats (TRs). Below the 2-D alignment (underneath the underlining) is a self-consensus and some expected conflicts (non-consensus) highlighted in yellow. **(c)** A subsequent *cinch-k* module cinches intra-TR <u>*k*</u>-mers as shown. In the consensus row, a new TR repeat pattern is revealed. **(d)** A later *cinch-d* module cinches the *<u>d</u>e novo* or inter-TR repeats revealed previously in the consensus row. Other important component serial modules of *maximal*, such as the critical *cyclelize* module, are described in the text and other figures (Fig. 6).

In our *maximal* MHA implementation, we use a nucleotide substitution matrix that gives ½ the maximum identity score to transition substitutions. We also use a sparse matrix approach for efficiency and in particular find that we only need to fill in values in a hemi-diagonal band. A bandwidth of ~200 bp is sufficient due to the known higher frequencies of tandem repeats (TRs) at increasingly lower *k*-mer sizes and their repeat number. Furthermore, the *maximal* preparation of sequences for MHA first involves construction of a self-MHA path box (Fig. 6b). In other words, the first part of alignment of sequences is to themselves. In various contexts, we refer to this procedure as MHA-preparation, self-MHA, and the “cinching” of 1-D lengths and/or 2-D widths into narrower 2-D widths.

In *maximal*, an initial computational module called *cinch-t* (t = tandem repeats) traverses a path unlike that of the trace-back strategies used in traditional GA. First, the *maximal* implementation of MHA completely dispenses with the need to evaluate at each intersection whether to do diagonal matching or horizontal or vertical path translations for insertions and deletions. Instead, the *cinch-t* module uses a cinch-path finding strategy that begins in the upper left-hand corner and proceeds column by column from left to right, and from top to bottom in each column never passing the diagonal (hence the efficiency of a hemi-diagonal sparse matrix, see Fig. 7a). This path will define the initial 2-D alignment. At column positions less than the bandwidth (the bandwidth defines the sparse matrix version of the path box), one begins at the first (top) row of each column, but after these initial columns one simply begins at the first intersection further below that is within the bandwidth. At each intersection with score *S*(*m*, *n*) > 0 for *n* > *m*, one evaluates whether the sum score of the diagonal beginning at that intersection and of length *k* = *n* – *m* surpasses a score threshold, which in our implementation is based on a *k*-mer dependent fraction of allowed transitions. We currently discard diagonals if they have a single non-transition mismatch, but different substitution strategies can be used.

#### Width cinching

If a *k*-mer diagonal is found to surpass threshold, it is “cinched”, by which we mean that the line is terminated with an inferred replication slip character represented by the forward slash (“/”). Then the second or more repeats are placed under the first repeat unit block in a series of rows, one per additional TR unit, as microparalogical alignment (see examples in Fig. 7b–d or previous figures). The slip character has no bearing on sequence length and is not meant to adjust spacing as the gap character does in GA. Instead the slip character merely conveys that the biological sequence continues on the next row below (see Fig. 7 b–d). Similarly, tick marks can be used to populate the upstream parts of rows that do not contain sequence merely as a visual guide for columnar alignment. These tick marks also do not have any bearing on alignment lengths.

In the end, the *cinch-t* module cinches the first tandem repeats of smallest *k*-mer size even if they are constituents of a larger unit TR block of length *l* > *k* but then only cinches the larger *l*-mer TR blocks after that and so on (compare the Fig. 7a path box to the *cinch-t* output in Fig. 7b). In a subsequent *maximal* module, called *cinch-k*, small TRs within the bigger *l*-mer repeats after the first unit *l*-mer block are then cinched like they were in the initial *l*-mer block (Fig. 7c). During the initial *cinch-t* pass, a modest amount of computation is also expended in considering slips located in the slip shadow represented by the window from *m*+1 to *n*–1 within the first unit block. We implement additional non-obvious modules made possible by this general strategy and describe these below (for one example, see Fig. 7d).

Implementation of MHA in the *maximal* program reveals key aspects of the nature of genetic sequence. Foremost is that such sequences are delocalized in two dimensions and are inherently non-monotonic. GA essentially is based on the assumption of intrinsic monotonicity of sequence. By monotonicity of a genetic sequence we mean that its positional homology has a 1-to-1 linear relationship with successive nucleotide subunits indexed in the 5´ to 3´ direction. For example, in a 1-D model of sequence, there is no ambiguity as to positional order. In a 2-D model of sequence, one index is required to specify column position *n* along a consensus sequence, while a second index is required to specify which specific paralogy block (row *m*) of a TR series.

If *m* and *n* are two positions in a 1-D string such that *m* is upstream of *n*, in a 2-D sequence *n* might well be located upstream of *m* due to its location in the upstream half of a downstream block of microparalogy. This is not meant as a superficial observation based only on the 2-D self-alignment, because this delocalization may really apply to ancestral positional homology and/or to different MHA-based cyclelizable “cinching” as explained below (also see Fig. 8). Delocalization is a natural consequence of allowing one-to-many microparalogy.

**Fig. 8.**
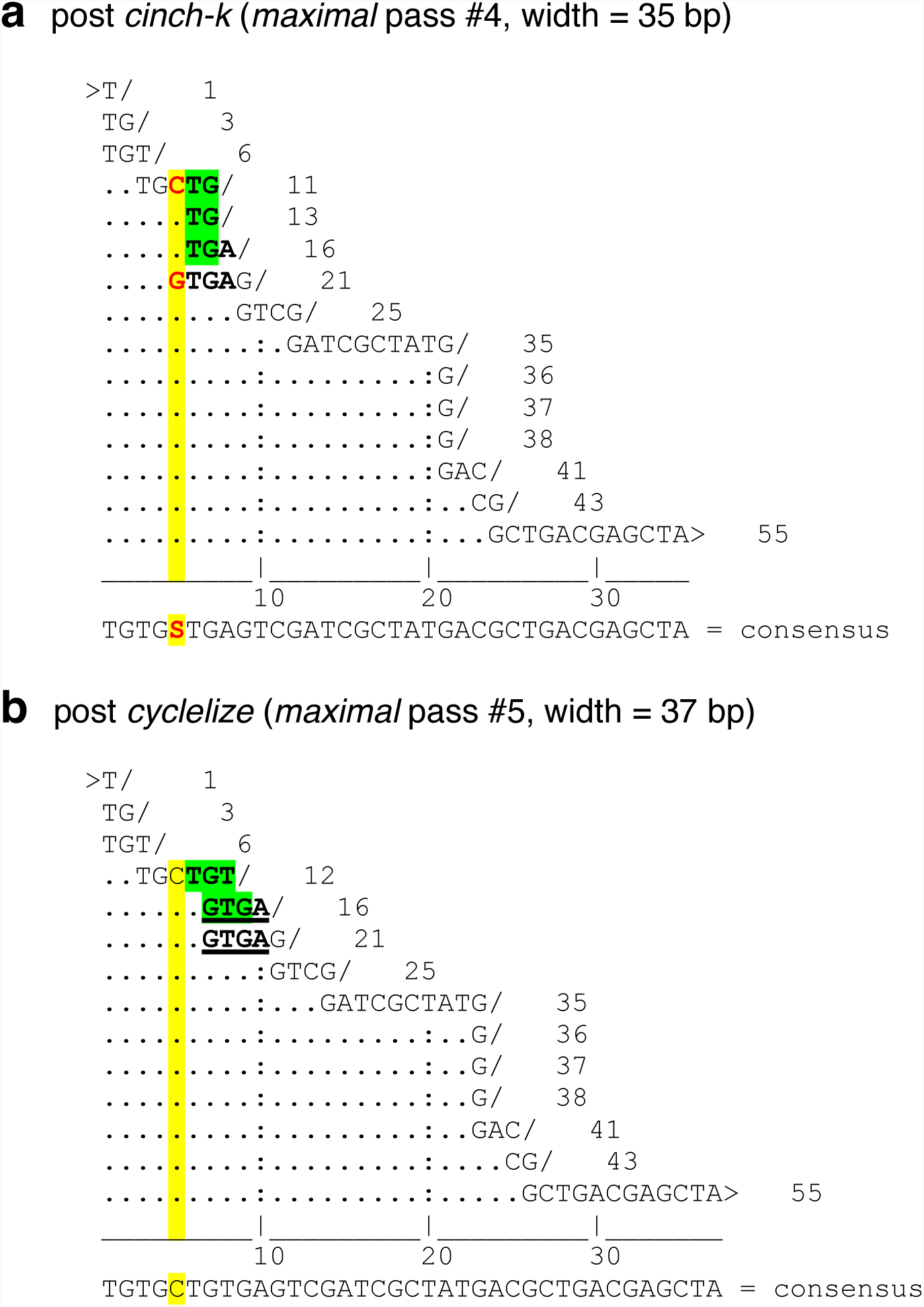
Cyclelizing overlapping and conflicting TRs. Shown are consecutive passes of a 2-D alignment by the *maximal* program in which a “cyclelizing” operation of a cycle sequence is required. Cycle sequences are defined as TRs that can be cinched in more than one frame. **(a)** Shown is a post *cinch-k* (pass #4) output revealing conflicting consensus (yellow highlighted column with red letters) due to the presence of overlapping repeats and because the cycling-frame of the first *k* = 2 repeat 5´-(TG)_3_ (highlighted in green) conflicts with the *k* = 4 repeat 5´-(GTAG)_2_. **(b)** This conflict is handled by the *cyclelize* module, which cycles the heptamer 5´-TGTGTG into 5´-T(GT)_2_G thereby rescuing the two repeats of the *k* = 4 TR (underlined). This resolves the previous non-consensus conflict, which was present in alignment column 5 (yellow column).

Replication slippage generates repetition that is increasingly unstable, leading to the generation of “micro-foam” microparalogy. Micro-foam sequence can be described as a complex pattern of repeats of repeats leading to private (lineage-specific), complexly homologous microparalogy. In a micro-foam model of sequence the concept of monotonic sequence position is untenable. For example, practical MHA display (*i.e*., capable of being graphed and represented 2-D dimensionally) must be able to “cyclelize” repeats. Cyclelizable tandem repeats must be “re-cinched” in a different repeat frame, say by first cinching 5´-**TGATTGATTGATTTACGATTTAC** into 5´-(**TGAT**)_3_**TTACGATTTAC** and then cyclelizing the initial repeat to allow cinching of an adjacent TR that overlaps with the last unit of the previous TR: 5´-**T**(**GATT**)_3_(<u>**GATT**</u>TAC)_2_ with the underlined sequence representing overlap with the third repeat of the tetramer. (This particular example sequence has an overall “width cinch ratio” of 8/23 or ~ 0.35, for example. Width cinch ratios [WCR] are likely to be characteristic of genomes in much the same way as indel rates and microsatellite repeat content can be rapidly divergent.) Whether the ancestral sequence contained the tetramer repeat starting with a **T** as in 5´-**TGAT** or with a **G** as in 5´-**GATT**, is not clear as it is unlikely for the edges of unstable TR repeats to have been perfectly conserved. Its instability during lineage divergence will have repeatedly scrambled the monotonicity over this sequence. Thus, 2-D MHA is more accepting of the non-monotonicity of biological sequences, which evolve in part by replication slippage, which is exacerbated in unconstrained microfoam.

In the *maximal* program, cyclelizing is handled by a *cyclelize* module and an example output is shown in Fig. 8. The current *maximal* version does true cyclelizing for *k* = 2, and “fudge-cyclelizing” for k > 2. Fudge-cyclelizing is an approximate solution to a cyclelizeable-resolvable microfoam knot, whereby *maximal* simply pushes the non-consensus rows to the right. We estimate that *k* = 2 cyclelizable knots are of comparable number to all other cyclelizable knots, but again cyclelizing is only required when resolving conflict between non-compatible overlapping TR patterns.

In implementing our particular approach to MHA, we found that the cinching of small *k*-mers after the *cinch-t* → *cinch-k* passes, and in particular of homopolynucleotide (*k* = 1) runs, allows recognition of differently-sized tandem repeats in the consensus row. For example, imperfect repeats differing by expansions and contractions of an internal homopolymeric run in the repeat unit are perfect repeats in the consensus row. Because the blocks differ in length, they are not seen by the initial *cinch-t*, *cinch-k*, and *cinch-s* modules. In the *maximal* program, these TRs of non-uniform size are self-aligned by a *cinch-d* module, which thereby effectively cinches *<u>d</u>e novo* (newer) inter-TR repeats of repeats (Fig. 7d). Different *maximal* modules can be evaluated as to their effectiveness in cinching by their before-and-after WCR ratios. We find that *cinch-d* provides one of the lowest (most effective) cinching ratios after *cinch-t* and is consistent with a known role of repeat slippage engendering further instability.

Our MHA program called *maximal* (https://github.com/microfoam/maximal) was written in C to explore MHA preparation and alignment under different parameters and recognizes DNA, RNA, protein, and alphabetical strings. With DNA sequence detection, transition matching is allowed and N’s (and other IUPAC DNA consensus characters) are rendered as non-cinchable lower-case ‘n’ characters. With protein sequence detection, N’s (asparagine’s) can be cinched and this is important because N’s are secondary structure breakers enriched in intrisically-disordered protein domains characterized by microfoam enrichment at the nucleotide-level (Fuxreiter *et al*., 2008; Tóth-Ptróczy *et. al*., 2008).

As *maximal* is the first implementation of MHA preparation, we make no claims concerning MHA efficiency. We do claim however, that formally-correct MSA for biological sequences subject to replication slippage must be of the MHA-type and not of the GA-type as they have been. MHA-approaches will be essential for functional sequence alignment in regions that are not constrained by codon-based open reading frames, which make it more likely that a replication slippage mutation is deleterious. MHA may also prove essential to resolving genome misassembly and mapping artifact (Ananda *et al*., 2013; Rice *et al*., 2015).

Our complete release 1.0 version of *maximal* (development version v2.74) is composed of a series of modules in the following order: *cinch-t* → *cinch-l* → *cinch-k* → *cyclelize* → *cinch-s* → *cinch-d* → *relax-2D* → *recover-1D* (available at: github.com/microfoam/maximal). The *cinch-l* module handles long homopolymeric repeats (*k* = 1) of length 2*m* or more, where *m* is a mononucleotide wrap length (m = 10), for a proportioned graphical aesthetic. In earlier versions of *maximal* the initial *cinch-t* was allowed to optionally proceed all the way down to *k* = *n*–*m* = 1, but we found that it is easier to allow *cinch-k* to cinch mononucleotide runs (and these only if they were smaller than the long runs handled by *cinch-l*, which wraps them in sets of 10 bp or at an optioned wrap length). Every module after the *cinch-t* module operates on the 2-D MHA rather than on the scoring path box, which is used only by *cinch-t*. The *cinch-k* module loops down to *k* = 1 after starting from the largest possible *k*-mer, which is the minimum of either *k* = *w*/2, where *w* is the band width of the sparse matrix, or else *k* = *r*/2, where *r* is the distance remaining to the end of the row (*i.e*., the next recorded slip). Within each *k*-mer loop of the algorithm, *cinch-k* proceeds row by row cinching *k*-mers as they are found left to right, and skipping rows immediately when the line is too short to harbor one. The *cinch-d* also loops from large to small *k*, but this module only consults the self-consensus row, and upon finding a TR there checks for microfoam conflict were it to be cinched. The *cinch-s* module cinches single-line TRs that do not affect global MHA alignment and which were invisible to *cinch-k*. The optional *relax-2D* module relaxes homopolymeric runs that did not aid in the cinching of repeats in the previous *cinch-d* runs and provides for a more aesthetically-pleasing MHA that has fewer rows. However, *relax-2D* is optimally dispensed for the purposes of MSA. Last, the *recover-1D* module recovers a 1-D dimensional string from the last 2-D cinching module and aligns it to the original 1-D string as a check to make sure a sequence is not scrambled unintentionally during cinching.

The program *maximal* was developed with an internal randomly-seeded Fisher-Yates randomization option (Fisher and Yates, 1938, 1948). During code development, we used this option to run tens of thousands of scrambled sequences from an enhancer (*vnd* NEE), from a protein-coding exon (CDS portion of *vnd* exon 3), and these from several divergent species of *Drosophila*. This procedure aided in the identification of “tricksy” strings harboring non-obvious, extra-tricky, microfoam knots, whose solution guided program development (tricksy strings are available with the code base). For example, we discovered a need to record TR cinching patterns in the initial *cinch-t* pass for checking potential slip conflict in the *k*-mer TR shadow, defined as the region of the first unit repeat, which may have been cinched inopportunely by *cinch-t* before the second unit has been identified (see Fig. 9). We thus devised a storage array that records slip locations by their number of repeats (sliploc[*i*]) and by their *k*-mer size (sliploc_nmer[*i*]). For compactly visualizing the number of repeats and their unit *k*-mer sizes, we use a single character base 62 system. This base 62 system uses the digits 0–9, the uppercase letters (A–Z), and the lowercase letters (a–z) to represent numbers 0 to 61, much as the hexadecimal system uses 0–9 and A–F to represent 0 to 15 (see example in Fig. 4 b). (For the rare occurrences where the number of TR blocks or the *k*-mer size is > 62, we use the “!” exclamation mark). This is also coupled with a single character display row (sliploc_echoes[*i*]) that is modified in regions of overlapping TRs and is used to store record of reversed slips (see Fig. 4b). The *maximal* program can thus display 1-D slip distribution patterns graphically using this MHA base 62 system and this is an informative aid for interpreting maximal 2-D cinching at regions of intense microfoam (see Fig. 4).

**Fig. 9.**
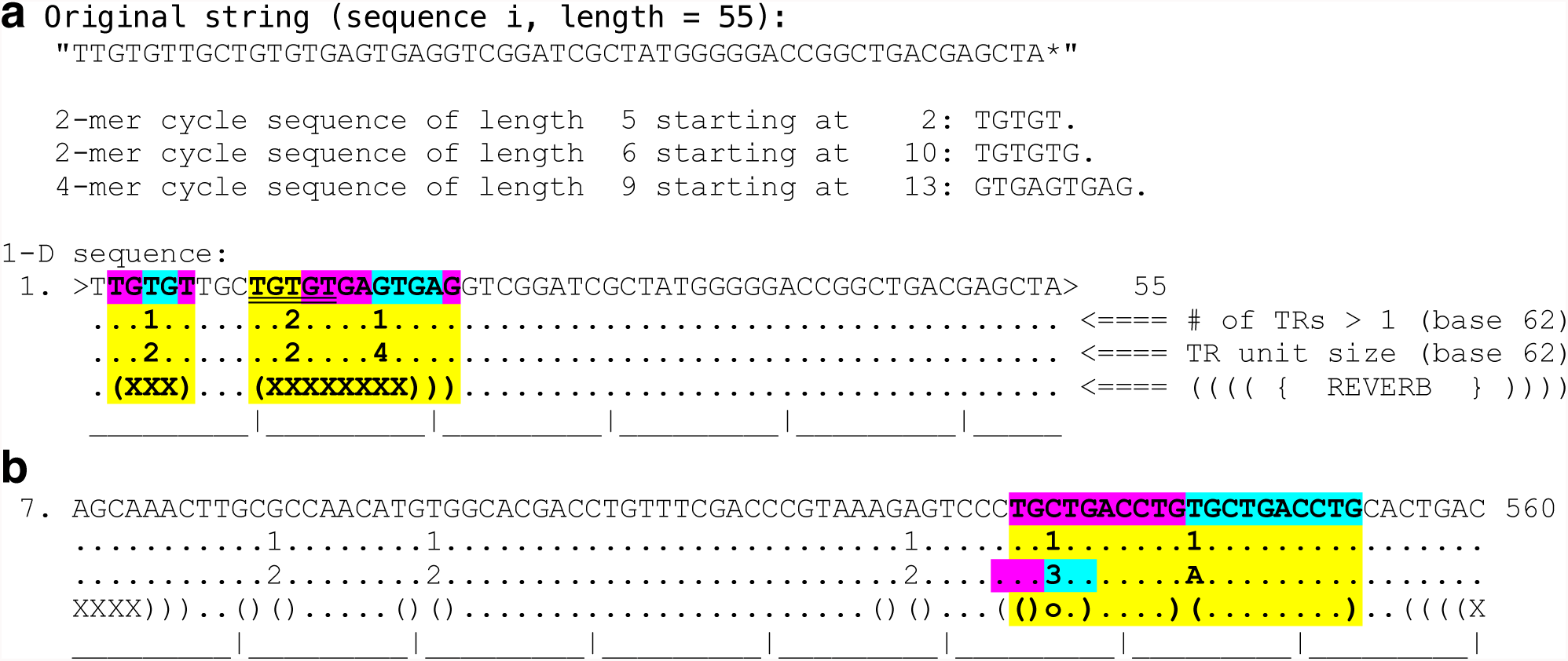
The program *maximal* displays 1-D slip location patterns compactly in base 62. **(a)** The use of a base 62 single character system (plus ‘!’ for *n* > 62) allows for compact display of the number of repeats in addition to the first unit block and the repeat’s *k*-mer size. A third row labeled “REVERB” marks the full extent of a cyclelizable sequence (see text for definition). Various characters are used for representing different types of overlap in the reverb or slip echoes row. For example, an ‘X’ marks that this location is simultaneously the beginning and end of a different TRs. In the first upstream example for the 5´-TGTGT sequence, this is due to the cyclelizable nature of the sequence. In other cases, it is due to separate overlapping repeats. This example fits in a single block so it begins and ends with the “>” symbol (block #1 of 1). **(b)** In this example of the *cinch-t* slip distribution in a window (block # 7 of 13 blocks) of the *D. melanogaster* NEE, we can see an example of the utility of the base 62 system at a TR of unit size *k* = A (10 in base 62). The ‘o’ character in the reverb line in the *k* = A slip shadow (the region of the first A-mer unit) marks a *k* = 3 TR that was dampened (reversed and subsequently ignored by other *maximal* modules) because its slip edge would have produced consensus conflict with the larger *k* = A repeat, which is a more productive cinch because it rescues a greater amount of microparalogy.

**Fig. 10.**
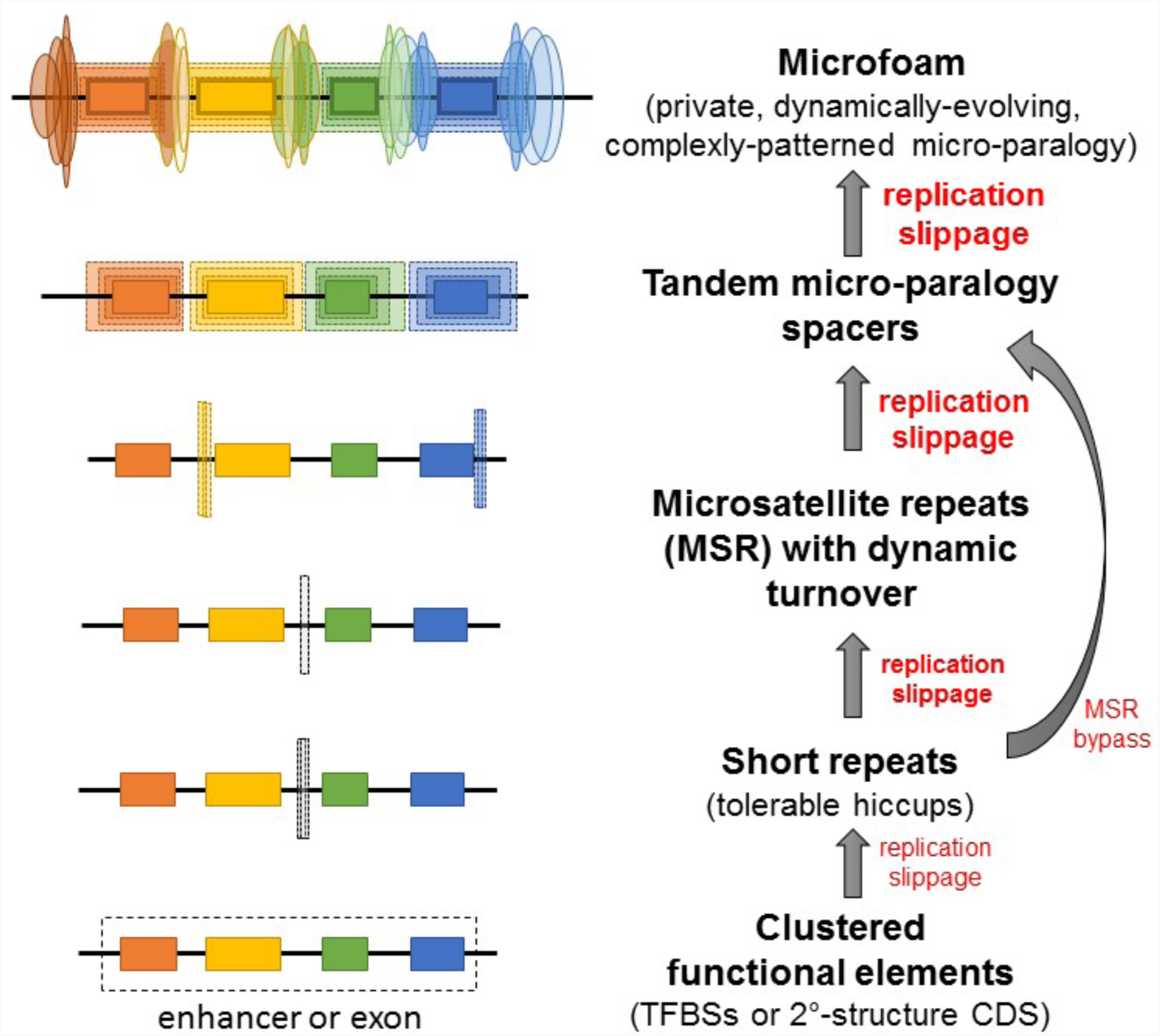
The microfoam model of sequence evolution. Shown is a model of the evolution of genetic modules encoding either *cis*-regulatory enhancer modules or protein-coding exons. In each such module, there is a finer sub-structure of elements (colored boxes). In a transcriptional enhancer, the sub-elements correspond to transcription factor binding sites. In a protein-coding exon, the sub-elements correspond to sequences typically encoding protein second-structural elements and other structural motifs additionally constrained by a translational reading frame (α-helices, β-strands, γ-turns, and protein-folding hydrophobic patches). As genetic sequences are prone to mutational changes related to the nature of the replication process, repeats produced by replication slippage (elongated dotted boxes) tends to accumulate with a bias in the spaces between the functional elements. This occasionally occurs on the edges of functional elements in which case the process can be described as echo sequences emanating from the functional elements. Furthermore, as the sub-elements are more resistant to evolutionary change, it is these sequences that frequently seed repeats into the spacer regions. Some of these repeats may evolve as tracts of perfect repeats known as microsatellite repeats (MSR) or tandem repeats (TR). However, this evolutionary model is envisioned as involving a great deal of imperfect repeats. Both perfect and imperfect repeats are increasingly unstable with increasing repeat number leading to a greater amount of repeat content. After much time, the less functional spacer sequences tend to resemble the functional elements via complex microparology, we refer to as microfoam (colored elongated ovals). Microfoam sequence delocalizes positional homology (the color of the ovals matching that of the adjacent rectangle elements represent paralogical relationship).

## Discussion

### Conclusion and significance

We conclude by addressing one possible misunderstanding about MHA relative to GA. It might be concluded that, like GA, the output of MHA is similarly fixated on an optimal alignment even if the MHA approach involved modeling a probabilistic distribution of such alignments. This is not exactly the case from the following thermodynamic perspective. MHA results in many fewer alignment columns than GA for the reason that microparalogy is rescued (see Table 1). In contrast, GA produces MSA’s with “width cinch ratios” > 1 because null characters are being inserted. In other words, with MHA there are many more ways to be homologously placed into the same column for the different letters of one sequence. The significance of this key difference between MHA and GA means that one can be homologous in many different ways inside an MHA alignment column. It is even possible for *different* (non-identical) versions of MHA-self alignment to be produced that are identical in their consensus sequence (and hence also their WCR). These *nearly-identical* MHAs would differ in the manner in which microparalogy is stacked in a subset of columns. By virtue of this fact, MHA relaxes the difficulty of placing characters into the correct alignment column because there are now more ways of being incorporated into that column. Thus, many different MHAs of the same sequence or set of sequences, all with identical consensus sequences, can be optimally and identically correct at the level of microhomology *sensu lato*. Furthermore, any one of these different MHAs are equally correct at the level of microhomology *sensu lato*.

## Acknowledgements

The author thanks Dr. Bin He and Dr. Gary Gussin for commenting on earlier versions of this manuscript. The author also thanks the Developmental Studies Hybridoma Bank that is run for the NIH by the University of Iowa for supporting travel to the Genetics Society of America’s 59_th_ Annual Drosophila Research Conference to present these results.

